# Rhizosphere-enriched microbes as a pool to design synthetic communities for reproducible beneficial outputs

**DOI:** 10.1101/488064

**Authors:** Maria-Dimitra Tsolakidou, Ioannis A. Stringlis, Natalia Fanega-Sleziak, Stella Papageorgiou, Antria Tsalakou, Iakovos S. Pantelides

**Author notes:** Shared first authors (Maria-Dimitra Tsolakidou, Ioannis A. Stringlis). **Corresponding author:** Iakovos S. Pantelides, **Address:** Department of Agricultural Sciences, Biotechnology and Food Science, Cyprus University of Technology, 30 Arch. Kyprianos Str., 3036 Limassol, Cyprus., **E-mail:**.

## Abstract

Composts represent a sustainable way to suppress diseases and improve plant growth. Identification of compost-derived microbial communities enriched in the rhizosphere of plants and characterization of their traits, could facilitate the design of microbial synthetic communities (SynComs) that upon soil inoculation could yield consistent beneficial effects towards plants. Here, we characterized a collection of compost-derived bacteria, previously isolated from tomato rhizosphere, for *in vitro* antifungal activity against soil-borne fungal pathogens and for their potential to change growth parameters in *Arabidopsis*. We further assessed root-competitive traits in the dominant rhizospheric genus *Bacillus*. Certain isolated rhizobacteria displayed antifungal activity against the tested pathogens and affected growth of *Arabidopsis*, and Bacilli members possessed several enzymatic activities. Subsequently, we designed two SynComs with different composition and tested their effect on *Arabidopsis* and tomato growth and health. SynCom1, consisting of different bacterial genera, displayed negative effect on *Arabidopsis in vitro*, but promoted tomato growth in pots. SynCom2, consisting of Bacilli, didn’t affect *Arabidopsis* growth, enhanced tomato growth and suppressed Fusarium wilt symptoms. Overall, we found selection of compost-derived microbes with beneficial properties in the rhizosphere of tomato plants, and observed that application of SynComs on poor substrates can yield reproducible plant phenotypes.

## Introduction

The soil layer surrounding plant roots, the rhizosphere, hosts a mesmerizing diversity of microorganisms that are highly affected by plant activity and represents one of the most complex terrestrial ecosystems^1,2^. The microbial communities in the rhizosphere, collectively known as the microbiome, are highly diverse and contain members having neutral, deleterious or beneficial effects on plants^2,3^. Generally, the structure of the microbiome is determined by the chemical environment of the rhizospheric soil, with main drivers being the root exudates^1,3^, the soil organic matter^4^ and exogenously-applied organic amendments^5^. Increasing evidence suggests that plants actively recruit beneficial bacteria that can either promote plant growth and health or can actively repress proliferation of soil-borne pathogens^6,7^. Thus, the root-associated microbiome is a subset of the more diverse microbiome of the bulk soil^8^. Perturbations in the composition of the root-associated microbiome can affect plant phenotypes, thus understanding the processes determining the composition of the rhizospheric community but also how specific members of the microbiome can help the plant are of high importance.

A considerable amount of research has been devoted to the identification of individual beneficial microbes responsible for plant growth and/or health. Plant growth-promoting rhizobacteria (PGPR) are widely studied beneficial microorganisms because of their ability to enhance plant growth and yield of several economically important crops. PGPRs can stimulate plant growth via the production of phytohormones (e.g. auxins, cytokinins and ethylene)^9^, the enzymatic activity they display (e.g. aminocyclopropane-1-carboxylate (ACC) deaminase)^10^ or by facilitating nutrition, e.g. phosphate solubilization^11^. PGPRs can also protect plants against soil-borne plant pathogens by producing antibiotics, hydrogen cyanide (HCN), and siderophores^12,13^, or through the activity of fungal cell wall-degrading enzymes^14,15^. Root colonization by some rhizobacteria can also induce systemic defense responses (induced systemic resistance; ISR) in the plant, that are effective against various plant pathogens in the leaves^16^.

The beneficial role of the root microbiome on plant health becomes more evident in disease-suppressive soils and organic amendments, in which disease will not develop even in the presence of a pathogen and a susceptible host^17^. Composts from different sources are the most widely-studied organic amendments that display suppressive properties against soil-borne pathogens and promote plant growth^18^. The phenomenon of suppressiveness is attributed to the microbial activity since it is abolished after sterilization or pasteurization^18,19^. A substantial number of bacterial, oomycete and fungal pathogens, including *Fusarium* and *Verticillium* spp., are targets of disease suppression^18^. Despite the extensive literature demonstrating the suppressiveness of composts, practical application is still limited due to the inconsistency of results and the unpredictable outcome in different pathosystems^20,21^.

The improvement of microbial isolation techniques allowed the identification of several microbial genera that are beneficial for plants^22,23^. Species belonging to *Bacillus, Enterobacter, Pseudomonas, Streptomyces* and *Trichoderma* are really active in the root environment and have the potential for disease control and plant-growth promotion^18^. However, to determine the contribution of individual strains from complex microbial communities to plant health is a daunting task. Various *in vitro* screening techniques are employed to identify microbes that could potentially improve plant growth and health; yet none of these screening methods guarantee plant-associated phenotypes^24^. Also, binary mutualistic interactions between plants and microbes are commonly applied for the identification of macroscopically visible beneficial outcomes but only a limited number have been applied successfully in practice^25^. The aforementioned suggest that these *in vitro* assays fail to predict *in vivo* effects as they cannot capture all variables of nature’s complexity.

In this study, we build on our previous work where we demonstrated that the activity of the microorganisms contained in a compost promoted plant growth and enhanced the protection of tomato plants against *Verticillium dahliae* (*Vd*) and *Fusarium oxysporum* f. sp *lycopersici* (*Fox*). Community-level analysis of the culturable microbiome revealed differential distribution of the microbial communities between the compost and the rhizosphere, suggesting that plants recruit beneficial soil microorganisms for their own benefit^26^. The aims of the present study are: (a) to assess if bacterial genera enriched in the rhizosphere of tomato plants have antifungal potential and can improve plant fitness; (b) to test whether the composition of synthetic communities (SynComs) can affect growth and defense outputs; and (c) to use simplified SynComs in sterile peat-based growth substrate and reproduce the suppressive and growth promotion properties of the compost on tomato plants. Using this approach, we provide with evidence that rhizosphere-enriched microbes can be pools for the construction of beneficial SynComs. Inoculation of an organic amendment lacking phytobeneficial properties with such SynComs can produce a growth substrate with predictable and consistent growth-promoting and disease suppressive properties.

## Results

### Production of antifungal and growth-modulating secondary metabolites is widespread among rhizosphere-enriched bacteria

To understand whether the previously observed disease suppressiveness of soil-borne pathogens^26^ is attributed to specific microbes selected in tomato rhizosphere, we tested in dual-culture assays the antagonistic activity of the rhizobacterial strains against *Vd* and *Fox* (Fig. 1). Initially, all 143 isolated strains were evaluated for direct antagonism through the production of diffusible secondary metabolites (DSM). A high percentage of these (32.9%, 47 strains) suppressed *Vd* growth whereas only 9 strains (6.3%) led to significant inhibition of *Fox* growth (Fig. 1a). Suppression of *Fox* ranged from 45.3% to 56.7%, while inhibition rates against *Vd* ranged from 3.3% to 51.3%. All the *Enterobacter cloacae* strains and strains *Bacillus licheniformis* (3Ba9, 3Ba17), *Bacillus subtilis* (1Ba19), *Chryseobacterium* sp. (2Ba19), *Ochrobactrum* sp. (2Ba57) and *Stenotrophomonas maltophilia* (2Ba31) were found to inhibit the growth of both *Vd* and *Fox*. From all the *Bacillus* strains tested, a relatively high percentage (42.6%, 20 isolates) could suppress mycelial growth of *Vd*. Next, we selected the 47 strains out of the 143 total isolates, causing significant *Vd* and *Fox* growth inhibition and tested their ability to suppress fungal growth through the production of VOCs (Fig. 1b). We observed a marked suppression of *Vd* by *Acidovorax* sp., that inhibited fungal growth by 55.3%, while 16 more isolates inhibited *Vd* growth more than 10%, with inhibition ranging from 10.3 to 28.4%. Twenty-eight isolates exhibited moderate inhibition, ranging from 0 to 9.6%, whereas strains *Microbacterium foliorum* (2Ba37) and *B. licheniformis* (3Ba44) slightly promoted *Vd* fungal growth by 2.5 and 1.5%, respectively. Regarding *Fox*, 18 strains led to pathogen inhibition ranging from 10.2 to 23%, whereas 21 strains showed less than 10% suppression. Seven *Bacillus* (1Ba2, 1Ba18, 1Ba43, 3Ba3, 3Ba30, 3Ba44, 3Ba45) and one *Microbacterium* (3Ba33) strains promoted *Fox* growth but to a small extent (0.5 to 4.4%). Interestingly, only three *S. maltophilia* (1Ba16, 2Ba27, and 3Ba7), one *Ochrobactrum* (2Ba57), one *E. cloacae* (2Ba47), one *Acidovorax* (1Ba38) and one *B. megaterium* (1Ba20) strains could suppress fungal growth of both pathogens more than 10% (Fig. 1b). This first experiment suggests that compost-derived bacteria isolated from tomato rhizosphere have the potential to inhibit fungal growth, mostly through production of DSM and in a lesser extent through VOCs.

**Figure 1:**
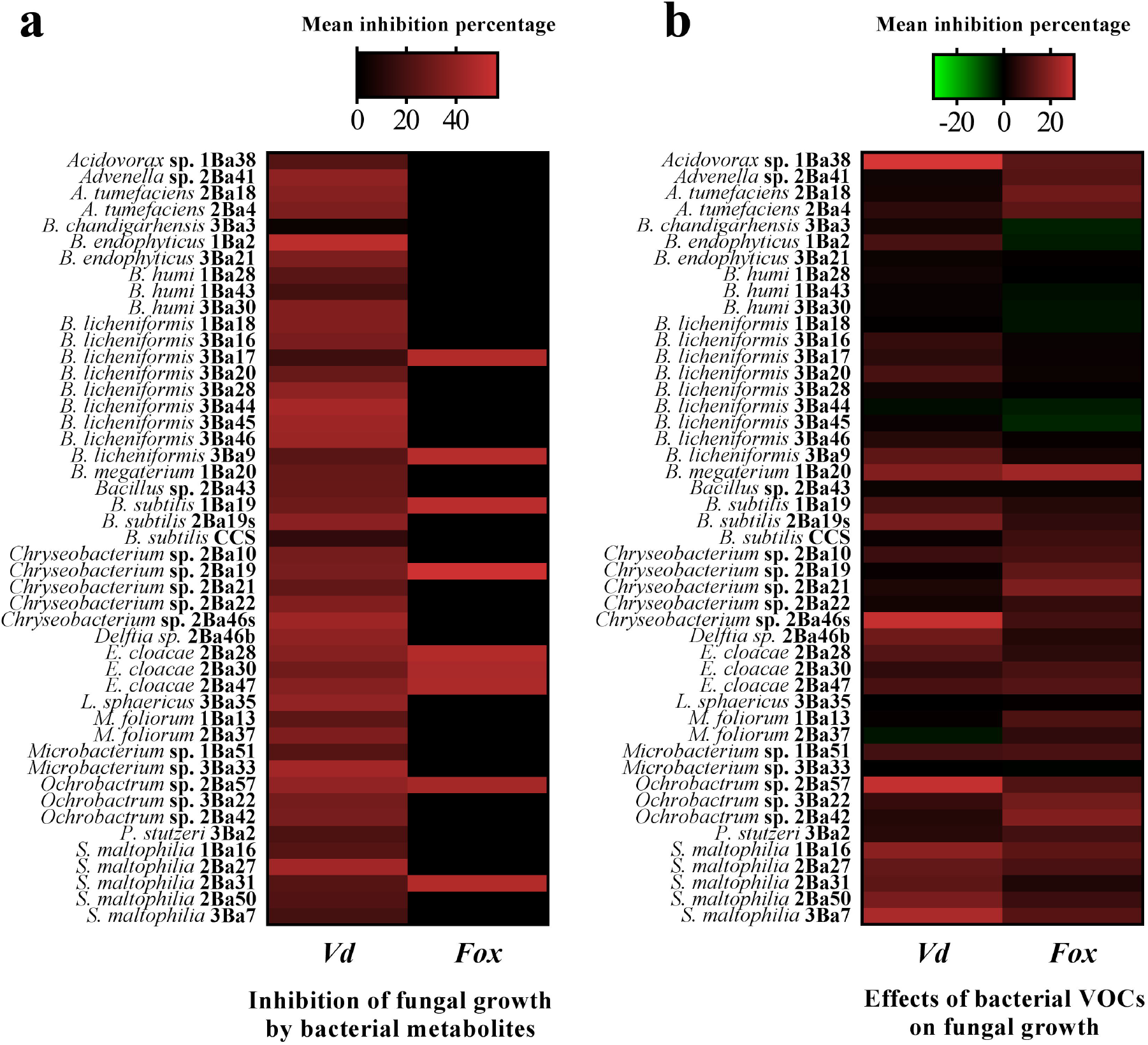
Antifungal potential of the rhizospheric microbial isolates. Heat maps illustrating the antifungal activity of (a) diffusible secondary metabolites and (b) volatile organic compounds (VOCs) of the studied bacterial isolates against the fungal pathogens *Verticillium dahliae* (*Vd*) and *Fusarium oxysporum* f. sp. *lycopersici* (*Fox*). Color of each box corresponds to the mean inhibition percentage recorded for each bacterial isolate as calculated from three and four replicate measurements, for diffusible secondary metabolites and VOCs respectively. Color scale ranges from green to red, with green, black and red representing negative values (promotion), baseline value (no effect) and negative values (inhibition), respectively. Measurements of antifungal activity were made 7 d after challenging *Vd* and *Fox* with the diffusible secondary metabolites or VOCs produced by the bacterial isolates.

Previously, the compost-derived and rhizosphere-enriched isolates were not only associated with disease suppressiveness, but also with plant growth promotion^26^. Therefore, the 47 strains displaying an inhibitory effect on *Vd* and *Fox* mycelial growth *in vitro* (Fig. 1) were tested for their ability to promote plant growth *in vitro*. Seven-d-old *Arabidopsis* seedlings were exposed to each bacterial strain and shoot biomass, primary root length and number of lateral roots were determined after 8 d of co-cultivation. Eleven out of 47 isolates significantly increased shoot fresh weight, 6 increased primary root length and 7 increased the number of lateral roots compared to the control (Fig. 2; green-colored boxes). A number of *S. maltophilia*, *E. cloacae*, *Delftia* sp., *Agrobacterium tumefaciens, Bacillus humi* and *B. licheniformis* strains led to significant increase in plant shoot fresh weight, whereas the remaining strains did not increase shoot biomass (Fig. 2a). Three *S. maltophilia* strains, *Delftia* sp., one *Chryseobacterium* strain and *Lysinibacillus sphaericus* significantly increased primary root length (Fig. 2b; green boxes), whereas *E. cloacae* 2Ba47 and 17 *Bacillus* strains showed negative effects on primary root length of seedlings (Fig. 2b; red boxes). Two *S. maltophilia* strains, two *Ochrobactrum* sp., one *E. cloacae*, one *Chryseobacterium* and *Advenella* sp. significantly increased the number of lateral roots (Fig. 2c; green boxes) whereas 12 *Bacillus* strains led to decreased number of lateral roots compared to the control (Fig. 2c; red boxes). The *B. megaterium* 1Ba20 induced negative effects on all plant growth parameters determined. These results indicate that the bacterial isolates when tested individually *in vitro* can have differential effects on plant growth, despite their isolation from the rhizosphere of plants showing growth promotion.

**Figure 2:**
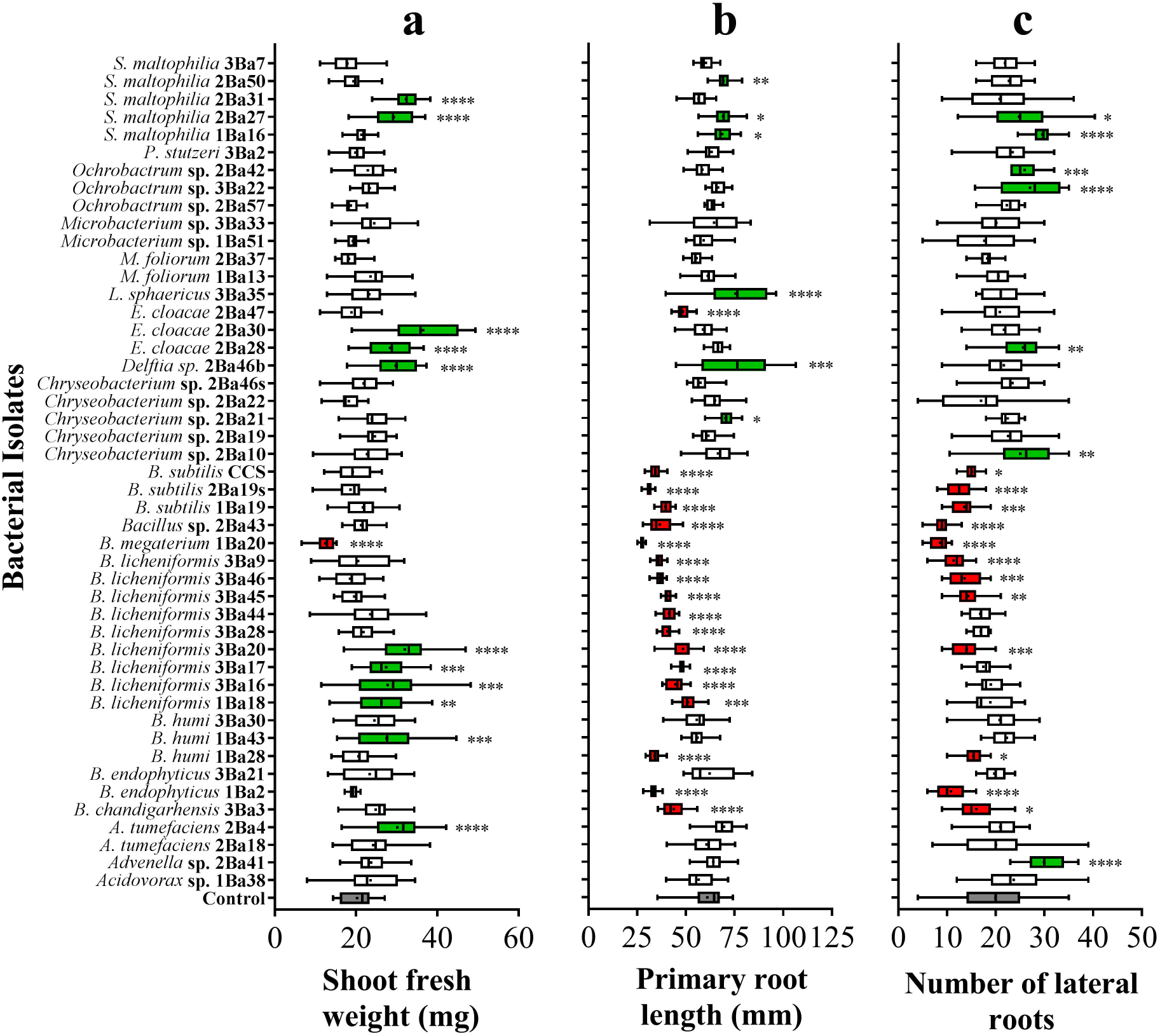
Effects of bacterial isolates on plant growth and root system architecture of *Arabidopsis* seedlings. Boxplots showing variation around the median for (a) shoot biomass production, (b) primary root length and (c) number of lateral roots, measured after 8 d of co-cultivation with each bacterial isolate (*n* = 21 plants). In each graph, grey box corresponds to control, red boxes to isolates with significant negative effect compared to control and green boxes to isolates displaying significant positive effect compared to control. White boxes represent strains with effect similar to control. Asterisks indicate statistically significant differences from control plants: \**P* < 0.05, \*\**P*□<□0.01, \*\*\**P*□<□0.001, \*\*\*\**P*□<□0.0001, oneway ANOVA, Dunnett’s test.

Another way to test if bacterial strains have the potential to affect plant physiology, is by evaluating their ability to produce/modify hormonal signals^48^. Auxin is one of the hormones closely related to plant growth and development^49^. Production of the auxin analog IAA was possible by 28 out of 47 strains (Fig. 3a). Four *Chryseobacterium*, 2 *Ochrobactrum*, one *S. maltophilia*, both *A. tumefaciens, Delftia* sp. and *Acidovorax* sp. strains produced IAA. A high percentage of the *Bacillus* strains assayed (85%, 17 strains out of 20) produced IAA, however at lower levels. Ethylene (ET) is a plant hormone that depending on its levels in plants can have either positive or negative effects on growth^50^. ET depletion is achieved by many PGPR strains through ACC deaminase enzymatic activity^10^. ACC deaminase activity was measured by determining the production of a-ketobutyrate from ACC. *Delftia* sp. 2Ba46b was the most efficient strain, among the 25 strains that exhibited ACC deaminase activity (Fig. 3b). All the *Delftia* sp., *Acidovorax* sp. *Advenella* sp., *Pseudomonas stutzeri, A. tumefaciens, Chryseobacterium* sp., *E. cloacae, Ochrobactrum* sp., *S. maltophilia* strains and 3 *Microbacterium* sp. strains were found positive to ACC deaminase activity. These assays show that a big proportion of the tested isolates have the potential to either synthesize hormonal analogs (IAA) or modify hormonal precursors (ACC), affecting as such plant growth.

**Figure 3:**
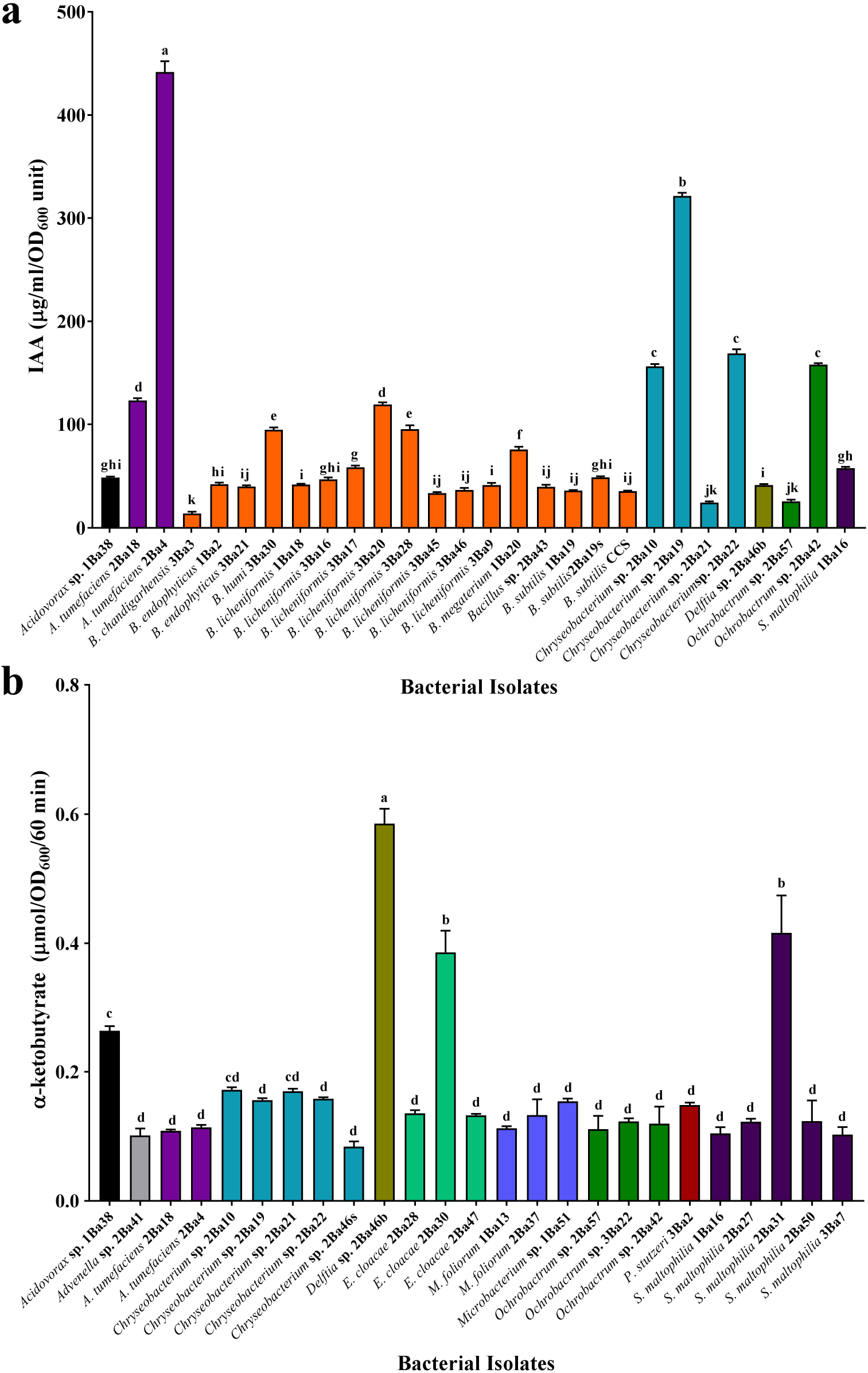
Production of IAA and cleavage of ethylene precursor ACC by bacterial isolates. Indole acetic acid (IAA) (a) and ACC deaminase activity (b) by bacterial isolates. Bars show the mean from three replicates of each isolate and are colored based on genus classification of the isolates. Error bars represent SE. Different letters indicate significant differences between the isolates (*P* < 0.05, oneway ANOVA, Tukey’s test).

### *Bacillus* rhizobacteria possess multifaceted beneficial characteristics

Enrichment of selected members of the microbiome in the rhizosphere can usually explain the beneficial effects conferred on plant fitness^16,51^. In our case, *Bacillus* isolates had increased representation among the rhizosphere strains (47 strains out of 143) accounting for 33% of the rhizospheric isolates (Supplementary Information Table S1). Hence Bacilli were selected to be characterized for an array of enzymatic activities and genetic markers associated with biological control activities. Microbial versatility in the rhizosphere and microbial biocontrol potential is usually linked to the production of hydrolytic enzymes (chitinases, glucanases, proteases, cellulases)^52,53^, while cyclic lipopeptides (CLPs) produced by microbes can inhibit growth of phytopathogens, facilitate root colonization and even stimulate host defense mechanisms^54^. Fifteen strains (71%) showed pectinase activity including all *B. licheniformis* and *B. subtilis* isolates; 8 strains (38%) exhibited cellulase activity; 6 strains (29%) showed proteolytic activity including all *B. subtilis* isolates; 10 strains (48%) showed chitinase activity including all *B. licheniformis* strains and eight strains (38%) showed phosphate solubilization potential including all *B. subtilis* strains (Fig. 4a). From these data, it becomes apparent that the isolated Bacilli constitute a pool of enzymatic activities.

**Figure 4:**
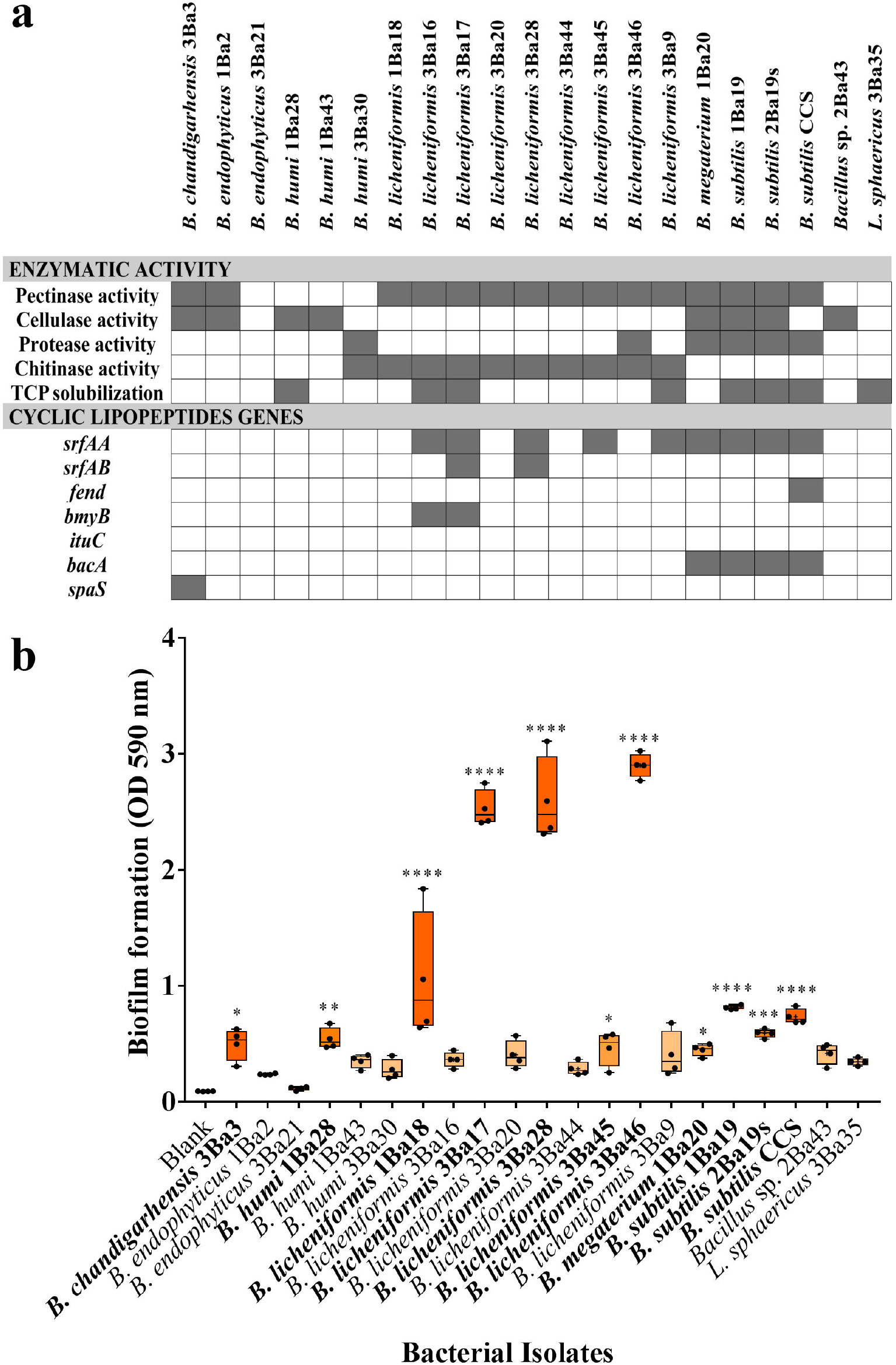
Enzymatic potential and biofilm formation of selected bacterial isolates. (a) Results of the different enzymatic activity assays and of the PCR targeting genes responsible for the production of cyclic lipopeptides in isolates of the genus *Bacillus*. Grey color indicates the presence and white color indicates the absence of the phenotype or gene in a bacterial isolate. TCP, Tricalcium phosphate [Ca_3_(PO_4_)_2_]; *srfAA*, surfactin synthetase subunit 1; *srfAB*, surfactin synthetase subunit 2; *fend*, fengycin synthetase; *bmyB*, bacillomycin synthetase; *ituC*, iturin A synthetase C; *bacA*, bacilysin biosynthesis protein; *spaS*, lantibiotic subtilin. (b) Boxplot of biofilm formation by each *Bacillus* isolate. Cells were cultured for 24 h in 96-well plates and the biofilm was quantified by crystal violet staining. Black dots represent the values of the 4 replicates per treatment. Asterisks indicate statistically significant differences from blank control: \**P* < 0.05, \*\**P*□<□0.01, \*\*\**P*□<□0.001, \*\*\*\**P*□<□J0.0001, one-way ANOVA, Dunnet’s test.

Except for general biocontrol activities, the 21 *Bacillus* strains were examined for the presence of CLP biosynthetic genes, since production of CLPs is linked not only to antimicrobial activity but also to biofilm formation^55^. Ten out of 21 strains had between one and three CLP biosynthetic genes. The most frequent gene marker was *srfAA* (9 positive strains), followed by *bacA* (4 positive strains). The gene markers *srfAB* and *bmyB* were found in two *B. licheniformis* strains (3Ba17, 3Ba28 and 3Ba16, 3Ba17, respectively), *fend* was found in *B. subtilis* CCS strain and *spaS* was detected in *B. chandigarhensis* (3Ba3) strain. The most frequent genotypes were *srfAA-bacA* found in all *B. subtilis* and *B. megaterium* strains and *srfAA-bmyB* found in two *B. licheniformis* (3Ba16, 3Ba17) strains. We further tested the *in vitro* ability of Bacilli strains to form biofilm (Fig. 4b). Eleven out of 21 strains (52%) showed significant biofilm formation with 4 *B. licheniformis* strains forming more biofilm than the remaining strains. Moreover, all *B. subtilis* strains were positive in biofilm formation and the same was observed for *B. chandigarhensis, B. megaterium* and *B. humi* 1Ba28 strains. Generally, the abovementioned data show that among Bacilli there are isolates with the potential to efficiently proliferate in the rhizosphere.

### Design of bacterial synthetic communities of different composition yields different outputs in *Arabidopsis* and tomato

In an effort to assess how the rhizosphere enrichment of diverse bacterial isolates from a compost can benefit the host plant^26^, we designed two simplified SynComs from the bacteria characterized in the current study. SynCom1 was composed of 25 strains, aiming to reflect the representation of the most abundant phylogenetic groups detected in the rhizosphere (Supplementary Information Table S3). SynCom2 was composed of 25 *Bacillus* strains (Supplementary Information Table S3), in order to test if the dominance of the Phylum Firmicutes in the rhizosphere of tomato plants (33%, Supplementary Information Table S1) can explain by itself the beneficial effects on plant fitness^26^. Initially, SynComs were tested *in vitro* for their effect on *Arabidopsis* growth, 20 d after root inoculation. Treatment of plants with SynCom1 caused adverse effects in plant growth and resulted in reduction of shoot and root fresh weight and root length (Fig. 5). In contrast to SynCom1, treatment of plants with SynCom2 did not affect shoot and root fresh weight, however primary root length was shorter compared to control plants (Fig. 5b). Root hairs’ mean length of SynCom2-treated plants was significantly longer compared to the controls, however the number of mature root hairs was not altered (Supplementary Information Fig. S1). These data show that while the SynComs contain microbes enriched in the rhizosphere of tomatoes showing growth promotion, *in vitro* assays with *Arabidopsis* cannot capture the complexity of the interaction between a community and a host that can support it.

**Figure 5:**
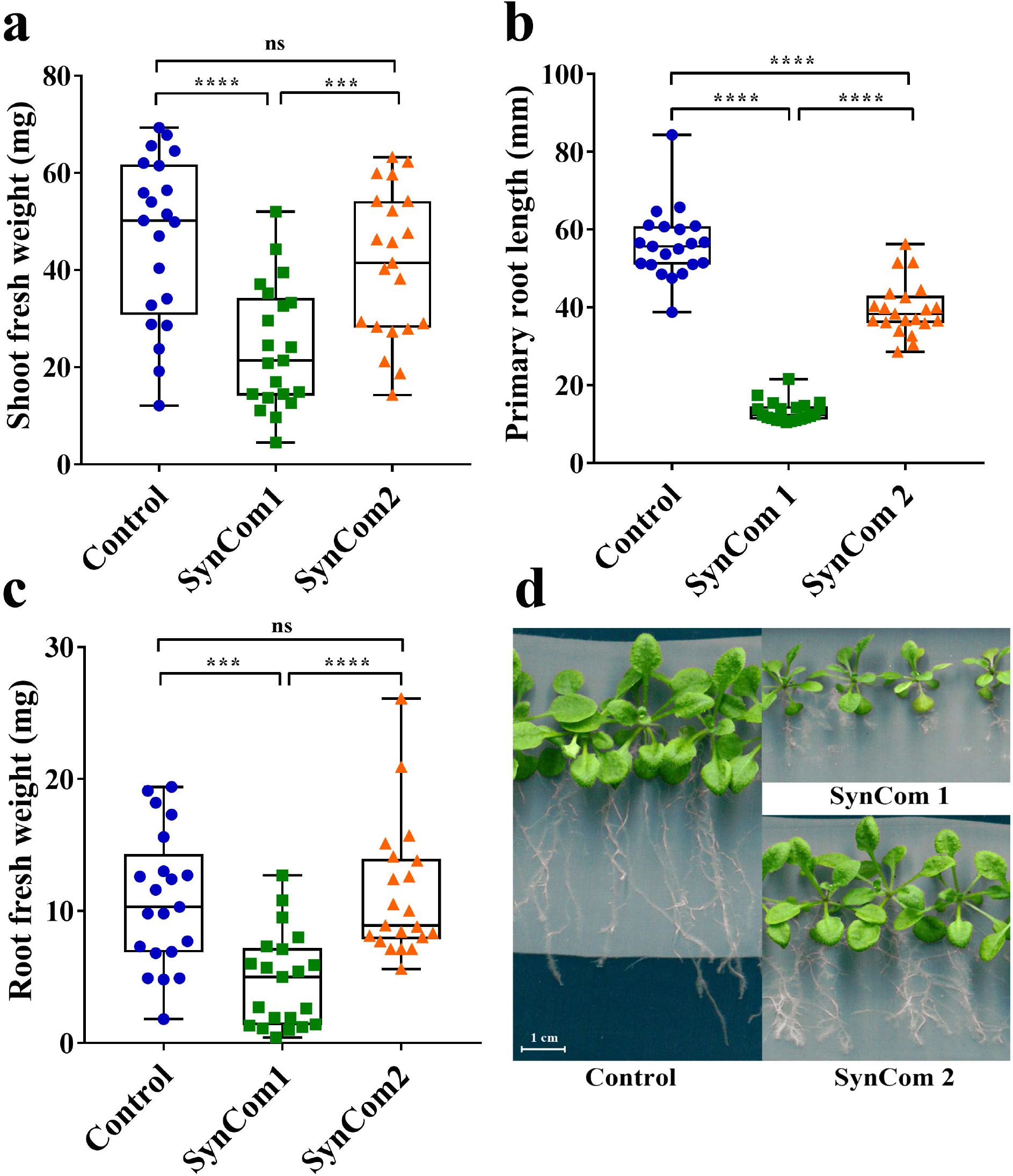
Effects of synthetic communities (SynComs) on *Arabidopsis* plant growth and root system architecture. Effect of bacterial synthetic communities on (a) shoot weight, (b) primary root length and (c) root weight of *Arabidopsis* plants (*n* = 21). Asterisks indicate statistically significant differences between treatments: \*\*\**P*□<□0.001, \*\*\*\**P*□<□0.0001, ns, not significant, one-way ANOVA, Tukey’s test. (d) Representative pictures of seedlings growing in control plates and plates with SynCom1 and SynCom2. Pictures were taken 20 d post co-cultivation.

To tackle the limitations of the *in vitro* experimentation, we tested the effect of the two SynComs on tomato plants in more pragmatic conditions performing pot experiments. In contrast to the *Arabidopsis* findings, in both SynCom treatments of tomato, the plants displayed increased growth as assessed by total leaf area, plant height and total number of leaves compared to the control plants (grown in sterile peat-based substrate) (Fig. 6). The effects were more striking in SynCom1-treated tomato plants compared to plants treated with SynCom2, as they showed greater growth in all parameters assayed (Fig. 6).

**Figure 6:**
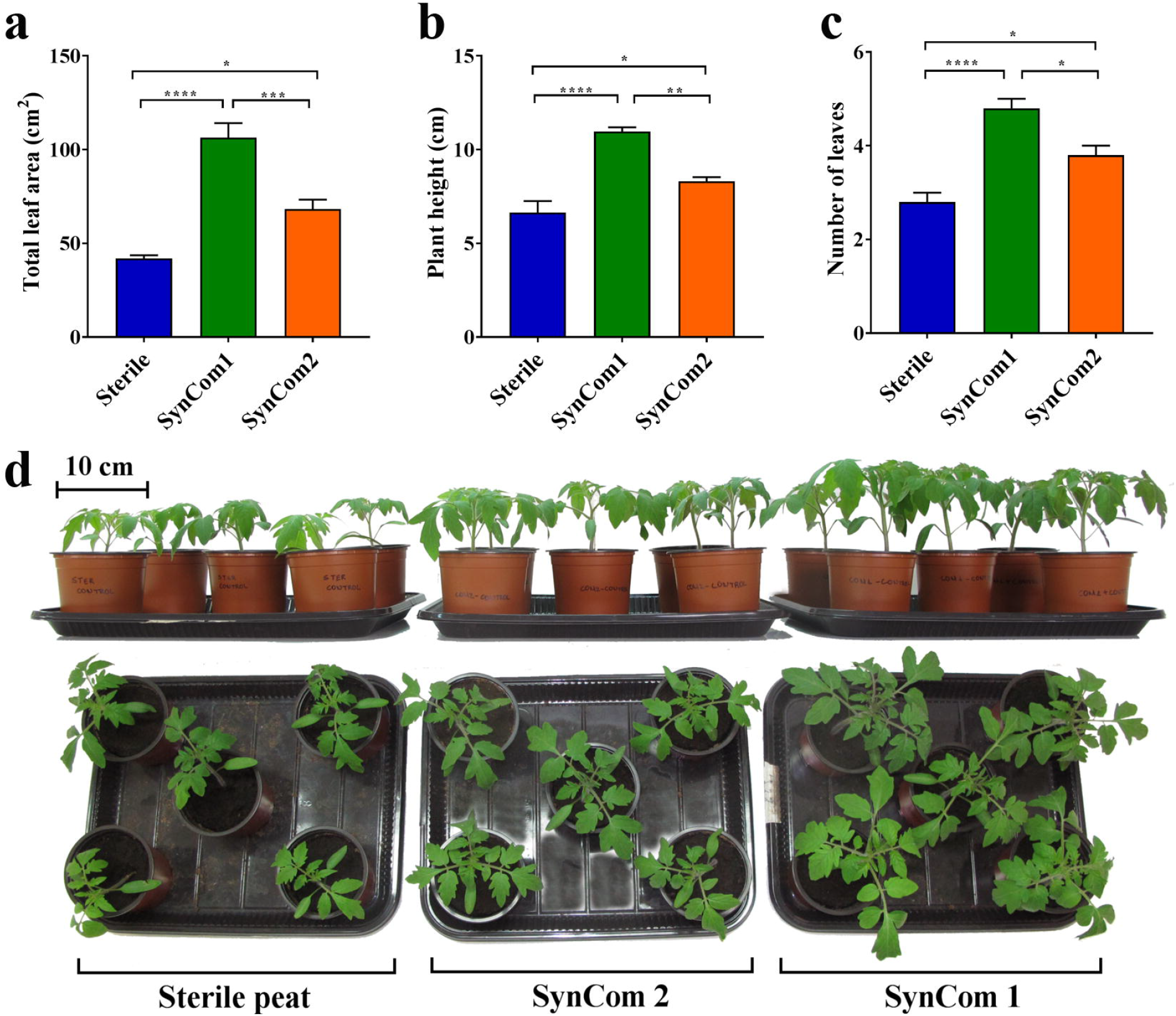
Beneficial effects of synthetic communities (SynComs) on tomato growth. SynComs-mediated effect on (a) leaf area (in cm^2^), (b) plant height (in cm) and (c) number of leaves. Data are means of 5 biological replicates and error bars indicate SE of mean. Asterisks indicate statistically significant differences: \**P* < 0.05, \*\**P*□<□0.01, \*\*\**P*□<□0.001, \*\*\*\**P*□<□0.0001, one-way ANOVA, Tukey’s test. (d) Representative pictures of tomato plants taken at 14 d after seedling emergence.

The members of the tested SynComs derive from a compost not only promoting tomato growth but also suppressing disease caused by soil-borne fungal pathogens^26^. Since both SynComs could benefit tomato growth, we then wondered if the tested SynComs could also reduce the symptoms caused by *Fox* after tomato infection. Interestingly, plants treated with SynCom2 showed less disease severity compared to the controls and SynCom1-treated plants (Fig. 7). The first *Fox* symptoms appeared in the form of wilting and yellowing especially on older leaves at 14 dpi and were recorder until 22 dpi (Fig. 7a). Disease symptoms progressed more rapidly in the control and SynCom1-treated plants, whereas SynCom2-treated plants showed less prominent symptoms and slower disease development (Fig. 7a, c). At 22 dpi, the disease severity in SynCom2-treated plants was 28.9% whereas in control and SynCom1-treated plants reached 58.4% and 48.7% respectively (Fig. 7b). The relative AUDPC values over 22 d of disease progress were higher in control (15.6%) and SynCom1-treated plants (15.9%) compared to SynCom2-treated plants (7.3%) (Fig. 7b). These data show that substrate inoculation with SynCom2 could reproduce the effect of the compost on disease suppressiveness. The ability of SynCom1 to promote growth in tomato but not to suppress disease, in the tested conditions, highlights the complexity behind naturally-observed growth promotion and further experimentation is required for its elucidation.

**Figure 7:**
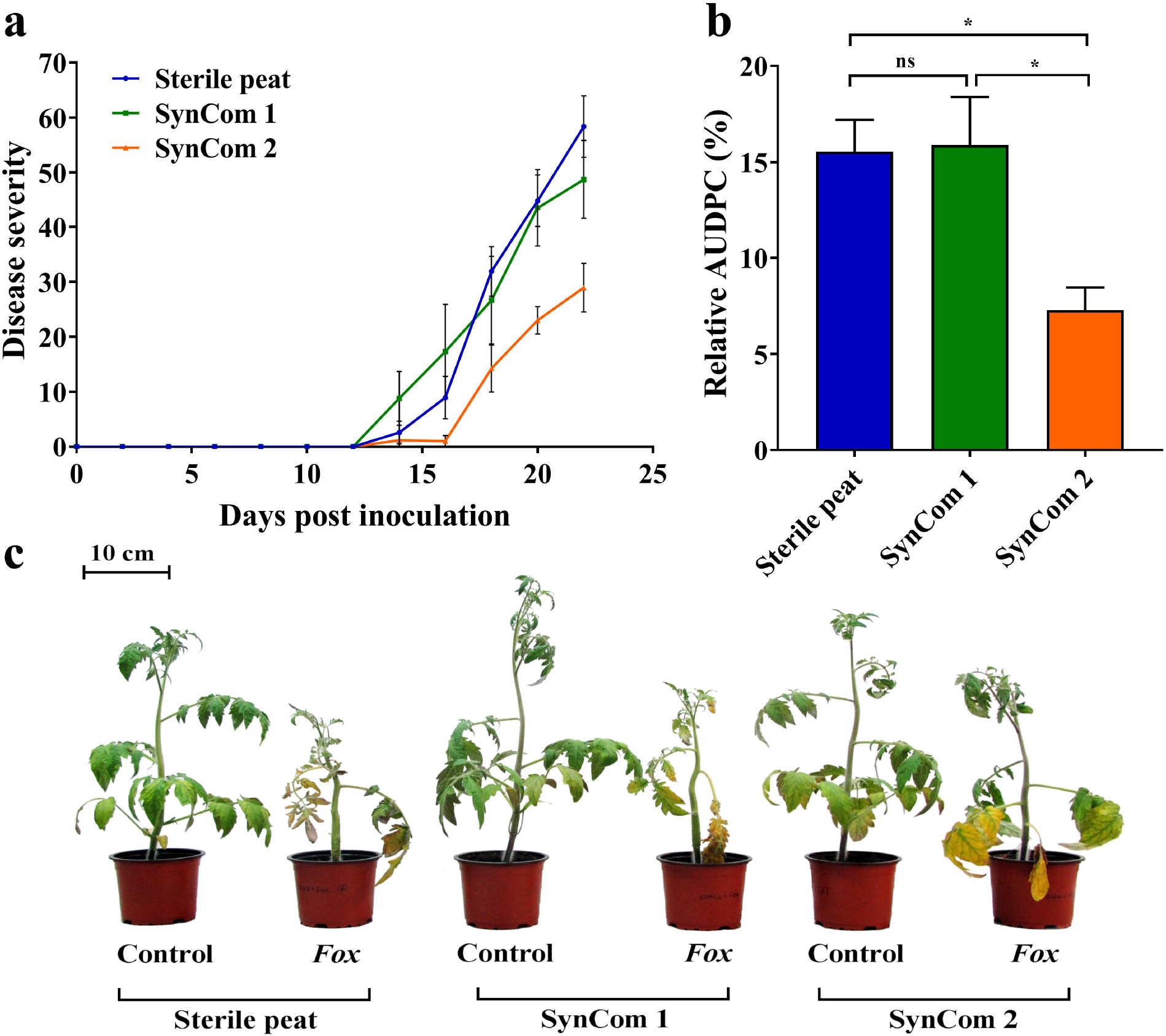
SynCom2 has a protective effect on tomato plants against *F. oxysporum* f. sp. *lycopersici*. (a) Disease progress over time and (b) amount of disease expressed as relative AUDPC. Error bars indicate the SE of mean (*n* = 8 replicates). Asterisks indicate statistically significant differences: \**P* < 0.05; ns,Jnot significant, one-way ANOVA, Tukey’s test. (c) Fusarium wilt symptoms on tomato plants grown in sterile peat (left), sterile peat inoculated with SynCom1 (middle) and with SynCom2 (right), at 22 d post inoculation.

## Discussion

Modern agriculture relies on the extensive use of chemicals to combat plant diseases and increase yields, which have nevertheless a negative impact on terrestrial ecosystems and human health. Therefore, the development and application of alternative and sustainable ways to deal with phytopathogens is a matter of urgency. In this respect, composts with suppressive properties are used to control soil-borne pathogens, that conventional methods, such as synthetic pesticides and resistant cultivars fail to control^18,21,56^. However practical application of composts in agriculture is limited due to their unpredictable and inconsistent growth-promoting and disease-suppressive properties^18^. Plants shape the structure of rhizosphere microbial communities by altering the chemical environment of the rhizosphere through root exudation^1^. Exudates can stimulate the abundance of selected microbial groups with beneficial properties that can largely contribute in the fitness and longevity of plants^6,7^. Several studies using cultivation-based techniques allowed the identification of antagonistic bacteria and fungi that when proliferating in the rhizosphere promoted plant growth and protected host plants from pathogens. In several cases, such isolated strains have been used to fortify composts and improve their consistency to control diseases^57,58^. Herein, we extend on our previous work by evaluating the antifungal and growth promotion potential of the bacteria enriched in the rhizosphere of tomato plants^26^, and by designing synthetic communities we aim to understand how the composition of microbial consortia can explain the enhanced growth and protection observed against the fungal wilt pathogens of tomato.

Evaluation of the bacterial collection enriched in the rhizosphere of tomato plants showed that strains from several genera displayed multiple plant-beneficial characteristics. A substantial number of *Bacillus*, *S. maltophilia*, *E. cloacae*, *Ochrobactrum* and *Chryseobacterium* strains inhibited *Vd* and *Fox* growth by producing DSM and/or VOCs (Fig. 1). The ability of *Bacillus* species to inhibit *Vd* and *Fox* growth both *in vitro* and *in vivo* has been extensively reported in the literature^59–61^. Similarly, antagonistic activity of *E. cloacae*, certain *S. maltophilia* and *Chryseobacterium* strains against *Fox*, *Vd* or both pathogens has been reported in *in vitro*, greenhouse and field trials^62–67^ whereas some strains belonging to *Ochrobactrum* suppressed the incidence of Fusarium wilt in greenhouse experiments^68,69^. Additionally, a large proportion of the tested isolates could produce IAA or displayed ACC deaminase activity *in vitro* (Fig. 3). Most *Bacillus* strains produced IAA, all *E. cloacae* exhibited ACC deaminase activity whereas strains of *Chryseobacterium*, *Ochrobactrum*, and *S. maltophilia* could produce both IAA and ACC deaminase. The potential of isolates belonging to the aforementioned genera or species to produce or modify hormonal signals is also well-documented ^68,70–72^.

Next, plant-bacterium binary-association experiments allowed us to assess the effect of each strain on plant growth, by using *in vitro*-grown *Arabidopsis* plants^23,42,43^. A substantial number of *Bacillus* strains increased *Arabidopsis* shoot fresh weight (Fig. 2a) and repressed primary root length and number of lateral roots (Fig. 2b, c). As shown in Fig. 3a, Bacilli are IAA producers and it was previously shown that microbes producing auxin or other molecules with auxin activity, can promote plant growth and affect root morphogenesis by inhibiting primary root elongation^73,74^. Most *E. cloacae* strains increased shoot fresh weight of *Arabidopsis* (Fig. 2), most likely via their ACC deaminase activity (Fig. 3b) by which bacteria can lower plant ethylene levels and stimulate plant growth^75^. Finally, certain *S. maltophilia* strains induced shoot growth, root length and number of lateral roots of *Arabidopsis* plants (Fig. 2) and these phenotypes could be attributed to IAA production and/or ACC deaminase activity (Fig. 3).

Release of various metabolites by the plants can attract beneficial microorganisms and enrichment of selected microbes in the rhizosphere has been shown to help plants deal with abiotic and biotic stresses^76,77^. More specifically, root associations with PGPR can protect plants from soil-borne pathogens by antagonistic mechanisms and/or activation of ISR^51,78^, and can help plants acquire more nutrients and grow bigger either by the production of phytohormones or the inactivation of environmental pollutants^79^. In our previous work, we found enrichment of Bacilli strains in the rhizosphere of tomato plants that displayed both increased growth and enhanced protection against soil-borne pathogens^26^. Therefore, we tested here whether individual Bacilli isolates retain properties that could benefit the plant. Apart from the antifungal activity and the growth-modulating compounds discussed earlier, we found that a large part of the *Bacillus* strains produced hydrolytic enzymes and possessed genes for CLPs biosynthesis (Fig. 4a). Previous studies have shown that hydrolytic enzymes, such as chitinases, proteases, cellulases, and glucanases, are involved in the antagonistic activity of *Bacillus* sp. and other biological control agents against fungal pathogens^14,15,80^. Production of antibiotics, including CLPs, is considered one of the most convincing properties of *Bacillus* spp. contributing to a broad-spectrum antimicrobial activity^81^. Besides their antimicrobial role, CLPs are also known to act as biosurfactants affecting the ecological fitness of the producing strains in terms of root colonization and biofilm formation^55^. Several works have shown that a variety of CLPs can stimulate host immune responses and trigger ISR in certain host plants^54,82^. In our study, the ability of the *Bacillus* isolates to form biofilm was verified *in vitro* (Fig. 4b). Previously, Chen *etal*^83^ suggested a positive correlation between the ability of *B. subtilis* to form robust biofilm and its biocontrol efficacy, and interestingly the formation of robust biofilms by *Bacillus* in defined medium was correlated with that on the plant roots. Another study also demonstrated the responsiveness of Bacilli towards plant cues, since root-associated biofilms of *B. subtilis* were triggered by exudates released from tomato roots^84^. Taken together, the enrichment of Bacilli in the rhizosphere of tomato and the overall ability of the tested *Bacillus* strains to produce an array of lytic enzymes, possess CLPs biosynthetic genes and form biofilms may explain the beneficial effects observed on the plants and the suppression of soil-borne pathogens.

The contribution of each microbe to a plant phenotype is a major challenge as many processes with regard to host fitness are accomplished by microbial synergies. While the majority of biological control efforts has focused on single organisms in the past, it is now possible to extend the analysis of plant protection to the full breadth of microbiota and develop microbiome-based biocontrol strategies^85^. Current research efforts focus on community rather than single-organism studies since the behavior of microorganisms in a community is different from that in pure cultures^86^. Studies have shown that single isolates with the ability to suppress fungal wilt pathogens under laboratory experiments often fail to induce disease suppression in the field due to low survival in the soil^87,88^. Other studies have linked species-richness of communities to higher resistance against pathogen invasions^89,90^. Therefore, practices that increase the size of the bacterial community may also increase the suppressiveness in soils and/or composts^89^.

Here, we constructed two SynComs using a bottom up approach by mixing selected bacterial strains that were exclusively recovered from tomato roots grown in a suppressive compost. The rationale behind these artificial assemblies was to achieve a SynCom composition reflecting the proportion of different Phyla in the culturable microbiome and in a next step to use those strains displaying *in vitro* antifungal properties against *Fox* and *Vd* (Fig. 1). We reasoned that inclusion of the most competitive bacterial isolates in terms of fungal growth inhibition and maintenance of this attribute in a community context, might result in a reduction of fungal wilt disease of tomato. For that, strain representation in SynCom1 was analogous to Phyla representation in tomato rhizosphere (Supplementary Information Table S3), in order to mimic the rhizosphere community features as closely as possible. SynCom2 contained only those *Bacillus* isolates that based on the *in vitro* pre-screening assays showed antifungal and plant growth-promoting potential. Application of SynComs on the roots of *Arabidopsis* and tomato plants in microcosm experiments produced differential outputs. SynCom1 was able to promote growth of tomato (Fig. 6) but compromised root length and fresh weight of *Arabidopsis* (Fig. 5). SynCom2 did not affect growth of *Arabidopsis* (Fig. 5) but promoted tomato growth (Fig. 6). Interestingly, SynCom1 that yielded the best growth-promoting effect on tomato, did not protect plants against *Fox*, whereas SynCom2 was able to reduce the severity of *Fusarium* wilt while displaying a moderate impact on plant growth (Figs. 6, 7). Our results suggest the critical involvement of *Bacillus* bacterial strains in the observed protection of tomato plants against *Fox*, while the combination of various different genera is required for the growth promotion of tomato. The phenotypic output of the *in vitro* experiments with SynComs and *Arabidopsis* plants were not consistent with the output of pot experiments with tomato plants suggesting that the effects produced my microbial consortia on host fitness are hard to predict only by *in vitro* experiments.

In conclusion, our results support the notion that growth promotion of plants and disease suppression are complex phenomena that are probably established by the activity of sophisticated microbial consortia^19,91^. The creation of synthetic ecosystems using microbial mixtures that can influence plant properties such as plant growth and disease resistance is an emerging research field that can lead to promising solutions for sustainable agricultural practices^43^. We propose that microbial synthetic communities can be used as compost inoculants to produce composts with desired characteristics e.g. predictive biocontrol of targeted pathogens. In that direction, further study is needed to understand how synthetic communities persevere in a compost and in the rhizosphere, what are the additive traits of a community compared to single strains and which host responses determine the outcome of the interaction.

## Methods

### Fungal species and bacterial strains

Race 1 isolate Fol004 of *F. oxysporum* f. sp. *lycopersici* (*Fox*)^27^ and race 1 isolate 70V of *V. dahliae* (*Vd*)^28^ were used. Studied bacterial strains (Supplementary Information Table S1) were isolated from the rhizosphere of tomato plants grown in a suppressive compost^26^. Both fungal and bacterial isolates were cryopreserved as an aqueous 20% glycerol suspension at -80°C. Fungal strains were activated on potato dextrose agar (PDA, Merck) at 25°C, while bacterial strains were grown on tryptic soy agar (TSA, BD) at 25°C.

### *In vitro* antifungal activity of bacterial isolates

The antagonistic activity of the diffusible secondary metabolites (DSM) of the bacterial isolates against *Vd* and *Fox* was determined using the *in vitro* dual-culture assay. PDA medium (20 ml) was poured into sterile Petri dish (90 mm diameter) and a 5 mm mycelial plug, taken from 6-d-old fungal culture, was placed on the center of the plate. Single bacterial colonies were streaked at ~1 cm from the plate’s rim and then plates were incubated for 7 d at 25°C. The inhibitory activity of bacterial isolates was expressed as the percent growth inhibition, compared to the control (plates inoculated with fungal plugs alone), according to the formula:

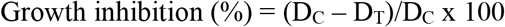

where, D_C_, diameter of control and D_T_, diameter of fungal colony in dual culture Each bacterial isolate was tested three times.

The effect of bacterial isolates VOCs on *Vd* and *Fox* growth was assessed using the bottoms of two Petri dishes. A 5-mm mycelium plug was added in one dish as above. Each bacterial isolate was inoculated into 25 ml tryptic soy broth (TSB, BD) in a 50 ml falcon tube and incubated for 48h at 25°C under agitation (180 rpm). A 100-μl bacterial culture (OD_600_=0.5) was spread on the second dish, containing 1x Murashige and Skoog medium (MS, Sigma-Aldrich) with 1.5% sucrose, 0.4% TSB and 1.5% agar^29^. The dish with the bacterial culture was inverted over the dish with the fungal mycelium plug, allowing physical separation between them and both dishes were sealed with Parafilm and incubated at 25°C. Petri dishes containing *Vd* or *Fox* exposed to a Petri dish containing only bacterial medium served as controls. Fungal growth inhibition was calculated as described above and each bacterial isolate was tested 4 times.

### *In vitro* plant growth assays

*Arabidopsis thaliana* Col-0 accession (hereafter *Arabidopsis*) was used in these assays. Seeds were surface sterilized^30^ and sown on 1x MS agar medium with 0.5% sucrose in square plates (120 × 120 × 17 mm). After 2 d of stratification at 4°C, plates were transferred in growth chamber (22°C; 16h light, 8h dark; light intensity 100 μmol m^−2^ s^−1^) and positioned vertically. Uniform 5-d-old seedlings were transferred to new square plates containing 1x MS agar with 0.5% sucrose. Each bacterial isolate was cultured on TSB and after 24–48 h of growth, cells were centrifuged at 2,600 *g* for 7 min at 4°C, washed twice in 10 mM MgSO_4_ and finally resuspended in 10 mM MgSO_4_. Ten μl of bacterial suspensions (final OD_600_=0.002) were spotted just below the root tip of each plant 2 d after transferring the seedlings to new plates. After 8 d of co-cultivation, shoot fresh weight, primary root length and number of lateral roots were measured as previously described^31^.

### Indole acetic acid (IAA) quantification assay

Production of IAA by the bacterial strains was measured in culture supernatants using Salkowski’s reagent^32^. Absorbance was measured at 535 nm with a Tecan Infinite M200 Pro plate reader (Tecan, Männedorf, Switzerland). The concentration of IAA was determined for each sample after comparison with a standard curve ranging between 5 and 50 μg ml^−1^.

### 1-aminocydopropane-l-carboxylate (ACC) deaminase activity

Each bacterial isolate was grown from mid-up to late-log phase in TSB in a shaking incubator (200 rpm, 25°C) and ACC deaminase activity was assessed by determining-ketobutyrate produced from ACC cleavage by ACC deaminase and comparing the absorbance at 535 nm to a standard curve of α-ketobutyrate ranging between 0.01 and 0.1 μmol^10^.

### Quantification of biofilm formation

Biofilm quantification was performed in 96-well plates as described before^33^ by staining biofilms with 0.1% (w/v) crystal violet (CV). CV concentration was determined by measuring the OD at 595 nm in a microtiter plate reader (Tecan). The samples were assayed in quadruplicate.

### Phosphate solubilization

Qualitative screening of phosphate-solubilizing bacterial isolates was performed on Pikovskaya agar medium medium containing 0.5% (w/v) Ca_3_(PO_4_)_2_. Bacterial isolates were streaked into Pikovskaya’s agar and phosphate solubilization was determined by the appearance of a clear halo around the colonies after 3 d of incubation at 28°C^34^.

### Determination of hydrolytic enzymes

Production of protease, chitinase, cellulose, and pectinase was detected on agar plates containing skim milk, colloidal chitin^35^, carboxy-methylcellulose and pectin, respectively^34,36,37^. Bacterial strains were inoculated on the respective agar plates and incubated at 28°C for 3-5 d. Production of clearing zone around the colonies indicated enzyme production. Zones of pectinase and cellulose were observed after staining with fresh Gram’s iodine solution^38,39^.

### Detection of cyclic lipopeptides (CLPs) genes

Total genomic DNA was extracted from bacterial isolates as described previously^26^. Primers used for the amplification of CLP biosynthesis genes^40,41^ are listed in Supplementary Information Table S2. PCR amplifications were carried out using Kapa Taq (Kapa biosystems) in a C100™ Thermal Cycler (Bio-Rad, Hercules, CA, USA) according to manufacturer’s recommendations (Supplementary Information Table S2).

### Synthetic community experiments Design of the synthetic communities

Two simplified (partially overlapping) synthetic communities (SynComs) of bacteria were designed (Supplementary Information Table S3) using representative rhizospheric bacteria isolated previously^26^. SynCom1 consisted of 25 isolates, selected to reflect the composition of the bacterial community, isolated from the rhizosphere of tomato plants grown in a suppressive compost (Supplementary Information Table S3). SynCom2 composed of 25 *Bacillus* isolates (from the same bacterial collection) was designed in order to test if the increased representation of the Bacilli compared to other genera in the rhizosphere of tomato plants (33%, Supplementary Information Table S1) can explain by itself the beneficial effects on plant fitness.

### Preparation of SynComs and treatments

Each bacterial isolate was grown for 48 h in TSB (180 rpm, 25°C). Cultures were rinsed with a 10 mM MgCl_2_ sterile solution and then centrifuged (2,600 *g*, 8 min). This process was repeated twice and cells were resuspended in 10 mM MgCl_2_ solution. The OD_600_ of each suspension was adjusted to 0.5 (~2.75 × 10^8^ cfu ml^−1^). SynComs were obtained by mixing the isolates in equal ratios. For the *Arabidopsis* experiments, each SynCom suspension was diluted to a final concentration of 10^5^ cfu ml^−1^^42,43^. *Arabidopsis* seedlings were inoculated by applying 10 μl of a bacterial suspension to the primary root of each seedling, immediately below the hypocotyl^44^. For SynComs tomato pot experiments, 2 ml of each SynCom suspension (OD_600_=0.5) was added to 50 ml sterile 10mM MgCl_2_ and mixed with 200 gr sterilized potting substrate (~2.75 × 10^6^ cells per gr substrate) directly before the sowing of surface sterilized tomato seeds.

### Plant growth conditions

*Arabidopsis* seeds were grown as described above. Uniform 6-d-old seedlings were transferred on 1x MS agar plates without sucrose, inoculated with a SynCom or MgCl_2_ (control) and were further grown for 20 d in a growth chamber (22°C; 16h light, 8h dark; light intensity 100 μmol m^−2^ s^−1^).

Tomato (*Solanum lycopersicum*) cv. Ailsa Craig seeds were surface-sterilized^45^ and sown into 10.5 cm diameter pots, each containing 200 gr sterilized peat-based potting substrate (Plantobalt substrate 2, Plantaflor)^26^, alone or with a SynCom. Plants were grown in a controlled growth chamber (25°C; 16h light, 8h dark; 65–70% RH; light intensity 450 μmol m^−2^ s^−1^ at pot height) and received watering individually every second day with equal volume of water.

### Experiments for the determination of plant morphological parameters

In the *Arabidopsis-SynComs* experiments, measurements of shoot and root fresh weight and primary root length were performed after 20 d of co-cultivation with the SynComs. For root hair length measurements, 5 individual roots per treatment were obtained and in each root the length of 27-34 root hairs was quantified by ImageJ (https://imagej.nih.gov/ij/). Totally, 307 root hairs were measured per treatment. Root hair density was determined as the average root hair number in a 1 mm^2^ root segment located 1 cm above the root tip. Root hair length and density were estimated using photographs taken by a binocular microscope (Olympus SZX16) with an attached digital camera (ColorView II, Olympus Soft Imaging System GmbH).

In the tomato-SynComs experiments, leaf area, plant height and number of leaves were determined 14 d after seedling emergence. For leaf area measurements, 5 plants were grown in sterile potting substrate alone or with a SynCom as described earlier, individual leaves were dissected and their area was measured with ImageJ software.

### Pathology studies

Pathogenicity experiments were performed on tomato plants at the four-leaf stage. Plants were grown in sterile potting substrate alone or with a SynCom as described above. A 5-d-old *Fox* liquid culture grown in sucrose sodium nitrate (SSN)^46^ in an orbital shaker (140 rpm) at 25°C, was passed through cheesecloth to remove mycelia. Conidial concentration was adjusted to 10^7^ conidia ml^−1^ and each seedling was inoculated with 10-ml suspension via root drenching. Control plants were mock-inoculated with 10 ml sterile distilled water. Disease severity was calculated by the number of leaves showing typical symptoms (wilting, yellowing, necrosis, stem collapse) as a percentage of the total number of leaves of each plant. Symptoms were periodically recorded for 22 d post inoculation (dpi) with the pathogen. Disease ratings were plotted over time to generate disease progress curves. The area under the disease progress curve (AUDPC) was calculated by the trapezoidal integration method^47^.

### Statistical analysis

Data were analyzed by one-way ANOVA (Tukey’s test with *P* ≤ 0.05 or Dunnett’s test with \**P* < 0.05, \*\**P*□<□0.01, \*\*\**P*□<□0.001, \*\*\*\**P*□<□0.0001), or two-way ANOVA (Sidak’s test with *P* ≤ 0.05) using GraphPad Prism 7 for Windows (GraphPad Software, La Jolla, CA, United States).

## Supporting information

## Data Availability

All data generated or analysed during this study are included in this published article (and its Supplementary Information files).

## Acknowledgements

This work was supported by the Cyprus University of Technology. The initial contribution of Michalis Aristeidou to this work is gratefully acknowledged.

## Author Contributions

IAS and ISP conceived and designed the research. MDT, NFS, SP, AT and ISP performed experiments. MDT, IAS and ISP analyzed and interpreted the data. IAS and ISP wrote the manuscript. MDT and IAS contributed equally. All authors have read and approved the final version of the manuscript.

## Additional Information

**Competing interests:** The authors declare no competing interests. No financial or personal relationship with other people or organizations exist that could inappropriately influence or bias this publication.

