## Supplementary material for "Rhizosphere-enriched microbes as a pool to design synthetic communities for reproducible beneficial outputs"

The following Supporting Information is available for this article:

**Fig. S1** Effect of synthetic community 2 (SynCom2) on *Arabidopsis* root hair formation.

**Table S1**. Rhizospheric isolates used in this study

**Table S2**. Oligonucleotide primers used to detect cyclic lipopeptide genes in *Bacillus* isolates.

**Table S3**. List of rhizospheric isolates composing the synthetic communities (SynComs) and their proportion (%) in the total isolated microbes.

**Fig. S1. Effects of synthetic community 2 (SynCom2) on *Arabidopsis* root hair formation.** (a) Root hair density of *Arabidopsis* seedlings after 20 d of co-cultivation with SynCom2. Root hair density is calculated as the average root hair number in the root segment located 1 cm above the root tip ($n$ = 5). (b) Average root hair length of *Arabidopsis* seedlings ($n$ = 5) growing on control and SynCom2-containing plates. Asterisks indicate statistically significant differences (***$P < 0.001$, Students t test). Representative images of *Arabidopsis* root hair formation in root tips (c and d) and in root segments located 1 cm above the root tip (e and f) following MgSO4 (control) and bacterial SynCom2 inoculation. Pictures were taken at 40X magnification. Scale lines represent 500 µm.

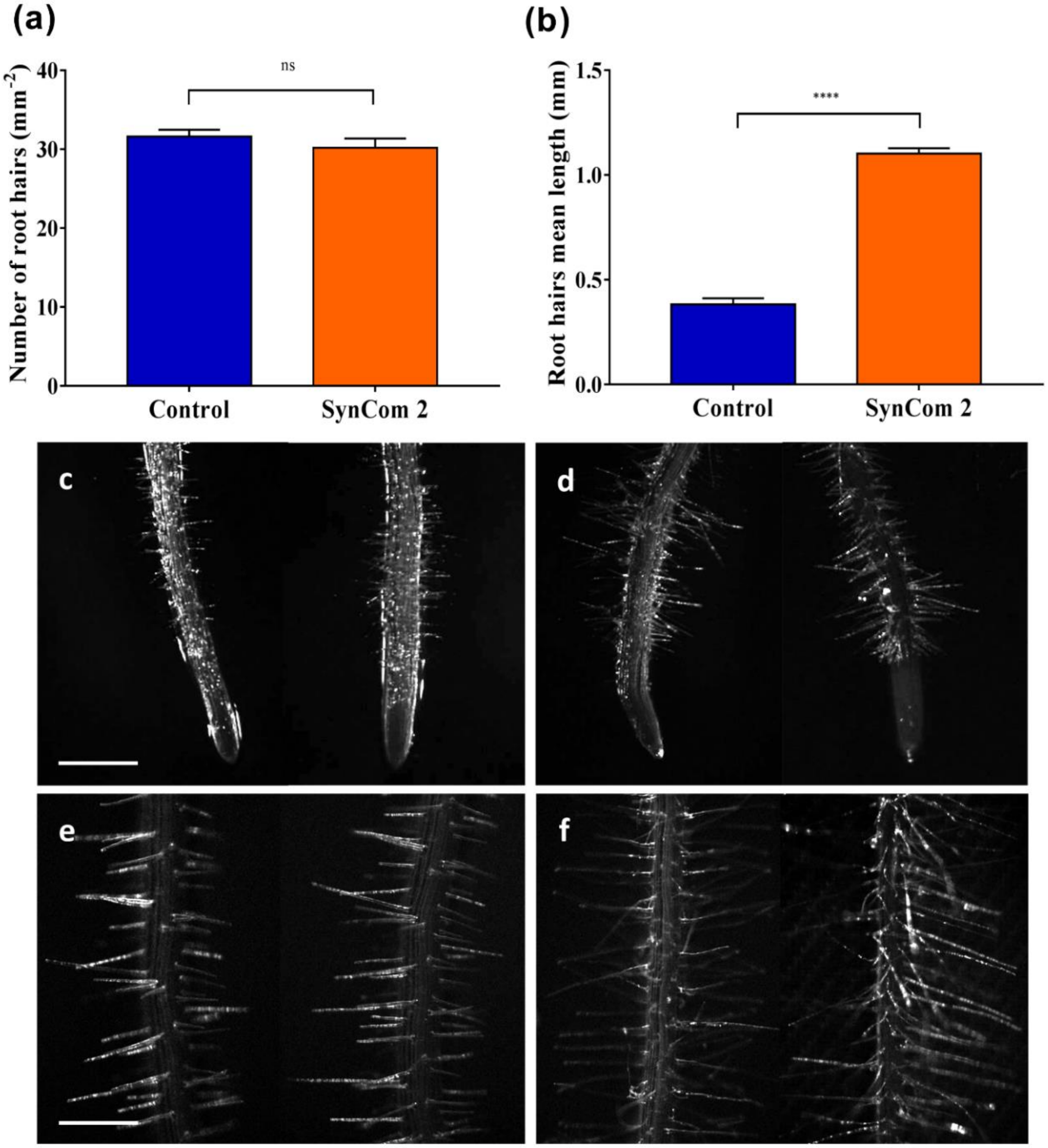

**Table S1**. **Rhizospheric isolates used in this study.**

This is a large Table in Excel and is submitted separately.

**Table S2**. **Oligonucleotide primers used to detect cyclic lipopeptide genes in *Bacillus* isolates.**

| Gene | Primer name | Primer Sequence (5’ → 3’) | Annealing temperature (°C) | Product size (bp) |
|---|---|---|---|---|
| Surfactin synthetase (*srfAA*) | SRFAF<br>SRFAR | TCGGGACAGGAAGACATCAT<br>CCACTCAAACGGATAATCCTGA | 58 | 201 |
| Surfactin synthetase (*srfAB*) | 110F<br>110R | GTTCTCGCAGTCCAGCAGAAG<br>GCCGAGCGTATCCGTACCGAG | 58 | 308 |
| Fengycin synthetase (*fenD*) | FNDF1<br>FNDR1 | CCTGCAGAAGGAGAAGTGAAG<br>TGCTCATCGTCTTCCGTTTC | 52 | 293 |
| Bacillomycin synthetase (*bmyB*) | BMBF2<br>BMBR2 | TGAAACAAAGGCATATGCTC<br>AAAAATGCATCTGCCGTTCC | 52 | 395 |
| Iturin synthetase (*ituC*) | ITUCF1<br>ITUCR3 | TTCACTTTTGATCTGGCGAT<br>CGTCCGGTACATTTTCAC | 52 | 575 |
| Iturin A synthetase C (*ituC*) | ITUCF<br>ITUCR | GGCTGCTGCAGATGCTTTAT<br>TCGCAGATAATCGCAGTGAG | 58 | 423 |
| Bacilysin biosynthesis protein (*bacA*) | BACF<br>BACR | CAGCTCATGGGAATGCTTTT<br>CTCGGTCCTGAAGGGACAAG | 58 | 498 |
| lantibiotic subtilin (*spaS*) | SPASF<br>SPASR | GGTTTGTTGGATGGAGCTGT<br>GCAAGGAGTCAGAGCAAGGT | 58 | 375 |

**Table S3. List of rhizospheric isolates composing the synthetic communities (SynComs) and their proportion (%) in the total isolated microbes.**

This is a large Table in Excel and is submitted separately.
